# Comparative analysis of protein abundance studies to quantify the *Saccharomyces cerevisiae* proteome

**DOI:** 10.1101/104919

**Authors:** Brandon Ho, Anastasia Baryshnikova, Grant W. Brown

## Abstract

Global gene expression and proteomics tools have allowed large-scale analyses of the transcriptome and proteome in eukaryotic cells. These tools have enabled studies of protein abundance changes that occur in cells under stress conditions, providing insight into regulatory programs required for cellular adaptation. While the proteome of yeast has been subjected to the most comprehensive analysis of any eukaryote, each of the existing datasets is separate and reported in different units. A comparison of all the available protein abundance data sets is key towards developing a complete understanding of the yeast proteome. We evaluated 19 quantitative proteomic analyses performed under normal and stress conditions and normalized and converted all measurements of protein abundance into absolute molecules per cell. Our analysis yields an estimate of the cellular abundance of 97% of the proteins in the yeast proteome, as well as an assessment of the variation in each abundance measurement. We evaluate the variance and sensitivity associated with different measurement methods. We find that C-terminal tagging of proteins, and the accompanying alterations to the 3’ untranslated regions of the tagged genes, has little effect on protein abundance. Finally, our normalization of diverse datasets facilitates comparisons of protein abundance remodeling of the proteome during cellular stresses.

---

Proteins are one of the primary functional units in biology. Protein levels within a cell directly influence rates of enzymatic reactions and protein-protein interactions. Protein concentration depends on the balance between several processes including transcription and processing of mRNA, translation, post-translational modifications, and protein degradation. The proteome within a cell is highly dynamic, and changes in response to different environmental conditions and stresses. Indeed, protein levels directly influence cellular processes and molecular phenotypes, contributing to the variation between individuals and populations (Wu et al. 2013).

Given the influence that changes in protein levels have on cellular phenotypes, reliable quantification of all proteins present is necessary for a complete understanding of the functions and processes that occur within a cell. The first analyses of protein abundance relied on measurements of gene expression, and due to the relative ease of measuring mRNA levels, protein abundance levels were inferred from global mRNA quantification by microarray technologies (Spellman et al. 1998; Lashkari et al. 1997). Since proteins are influenced by various post-transcriptional, translational, and degradation mechanisms, accurate measurements of protein concentration require direct measurements of the proteins themselves.

The most comprehensive proteome-wide abundance studies have been applied to the budding yeast model organism, *Saccharomyces cerevisiae*, whose proteome is currently estimated at 5858 proteins [*Saccharomy-ces* Genome Database, www.yeastgenome.org, accessed October 28, 2016]. Several methods for quantifying protein abundance have been employed, including tandem affinity purification (TAP) followed by immunoblot analysis, mass spectrometry (MS), and green fluorescent protein (GFP) tag–based methods. The generation of a yeast collection with each open reading frame (ORF) fused with a TAP tag allowed for one of the first global analyses of protein expression in yeast. Immunoblot and chemiluminescence detection of yeast extracts containing TAP-tagged proteins was performed, and absolute protein abundance determined by comparison to internal standards (Ghaemmaghami et al. 2003). Mass spectrometry further advanced global analysis of protein copy number in cells. The advantage of mass spectrometry-based approaches is that they do not require tagged proteins for quantification. Targeted proteomic approaches, including total ion signal, internal scaling, and selected reaction monitoring, have been used to quantify protein levels (Liebler & Zimmerman 2013). Collectively, TAP-immunoblot and mass spectrometry analyses of global protein expression provide highly sensitive measurements and 87% coverage of the yeast proteome.

While protein concentration largely influences cellular processes, protein localization has emerged as another important factor in protein functionality, prompting the construction of a yeast collection where each ORF is tagged with a GFP protein (Huh et al. 2003). This system allows simultaneous measurements of GFP-tagged protein intensity, as a proxy for protein abundance, and localization. The GFP strain collection has been extensively used to understand the cellular response to a variety of environmental conditions that include DNA damage, osmotic stress, starvation, and oxidative stress (Lee et al. 2007; Davidson et al. 2011; Tkach et al. 2012; Denervaud et al. 2013; Breker et al. 2013; Mazumder et al. 2013; Chong et al. 2015). GFP-based measurements of protein abundance typically correlate well with both TAP-immunoblot and mass spectrometry based approaches, suggesting that GFP fluorescence intensity is a reliable reporter for protein abundance (Torres et al. 2016). High-throughput microscopic studies have provided a comprehensive view of the plasticity of the yeast proteome under genetic and chemical perturbation (Tkach et al. 2012; Denervaud et al. 2013; Breker et al. 2013; Mazumder et al. 2013; Chong et al. 2015; Koh et al. 2015; Breker et al. 2014).

Existing protein abundance studies correlate well with one another, yet it remains difficult to derive reliable and accurate measurements of the abundance of any one protein, or of protein abundance across the proteome, from any one study. Only five existing data sets quantify protein abundance in molecules per cell (Ghaemmaghami et al. 2003; Kulak et al. 2014; Lu et al. 2007; Peng et al. 2012; Lawless et al. 2016), and no single study offers full coverage of the proteome. Proteome-scale abundance studies of the yeast proteome in the literature currently number nineteen (Ghaemmaghami et al. 2003; Newman et al. 2006; Lee et al. 2007; Lu et al. 2007; de Godoy et al. 2008; Davidson et al. 2011; Lee et al. 2011; Thakur et al. 2011; Nagaraj et al. 2012; Peng et al. 2012; Tkach et al. 2012; Breker et al. 2013; Denervaud et al. 2013; Mazumder et al. 2013; Webb et al. 2013; Kulak et al. 2014; Chong et al. 2015; Lawless et al. 2016; Yofe et al. 2016), providing an opportunity for comprehensive analysis of protein abundance in a eukaryotic cell. We describe such an analysis, incorporating all existing global studies of protein expression in yeast. We provide a single protein abundance estimate for each of 5702 proteins, covering 97% of the yeast proteome. We evaluate the protein concentration ranges that are most effectively measured by the existing methodologies. We find that two-thirds of the proteome is expressed within a narrow concentration range of 1000-5000 molecules per cell. Finally, we note that C-terminal fusion tags have only a modest effect on protein abundance.

## Materials and Methods

### Data Collection and Processing

We gathered 19 data sets from published studies measuring protein abundance across the yeast proteome, either reported in arbitrary units or in molecules per cell (Ghaemmaghami et al. 2003; Newman et al. 2006; Lee et al. 2007; Lu et al. 2007; de Godoy et al. 2008; Davidson et al. 2011; Lee et al. 2011; Thakur et al. 2011; Nagaraj et al. 2012; Peng et al. 2012; Tkach et al. 2012; Breker et al. 2013; Denervaud et al. 2013; Mazumder et al. 2013; Webb et al. 2013; Kulak et al. 2014; Chong et al. 2015; Lawless et al. 2016; Yofe et al. 2016). Throughout our analysis, we used designated codes to refer to each study (Table 1). Unperturbed measurements derived from (Chong et al. 2015) are the mean of the three technical replicates in their study, and for stress conditions the 160 minute data for hydroxy-urea and the 700 minute data for rapamycin were used. Measurements in unperturbed cells derived from (Denervaud et al. 2013) are the mean of all time points prior to their treatment condition. For (Peng et al. 2012), the average of all data was used. For (Webb et al. 2013), the average of the emPAI values from the three micro MudDPIT replicates was used.

**Table 1.** The nineteen protein abundance datasets considered

| Abbreviation | Reference | Type of Study | Detection | Abundance measure | Media | Temp | Growth phase |
| --- | --- | --- | --- | --- | --- | --- | --- |
| LU | Lu et al. 2007 [32] | Mass spectrometry | label-free spectral counting | absolute | YPD | 30°C | mid-log |
| PENG | Peng et al. 2012 [40] | Mass spectrometry | label-free spectral counting and ion volume based quantitation | absolute | Minimal |  | early log |
| KUL | Kulak et al. 2014 [25] | Mass spectrometry | label-free spectral counting | absolute | YPD | 30°C | mid-log |
| LAW | Lawless et al. 2016 [27] | Mass spectrometry | stable-isotope labeled internal standards and selected reaction monitoring | absolute | Minimal |  | chemostat |
| DGD | de Godoy et al. 2008 [12] | Mass spectrometry | SILAC and ion chromatogram based quantification | relative | Minimal |  | mid-log |
| LEE2 | Lee et al. 2011 [28] | Mass spectrometry | isobaric tagging and ion intensities | relative | YPD | 30°C | mid-log |
| THAK | Thakur et al. 2011 [48] | Mass spectrometry | summed peptide intensity | relative | Minimal |  | mid-log |
| NAG | Nagaraj et al. 2012 [36] | Mass spectrometry | spike-in SILAC | relative | YPD | 30°C | mid-log |
| WEB | Webb et al. 2013 [58] | Mass spectrometry | label-free spectral counting | relative | YPD | 30°C | mid-log |
| TKA | Tkach et al. 2012 [50] | GFP-microscopy | live cells; confocal | relative | Minimal | 30°C | mid-log |
| BRE | Breker et al. 2013 [5] | GFP-microscopy | live cells; confocal | relative | Minimal | 30°C | mid-log |
| DEN | Denervaud et al. 2013 [13] | GFP-microscopy | live cells; wide field | relative | Minimal | 30°C | steady-state |
| MAZ | Mazumder et al. 2013 [33] | GFP-microscopy | fixed cells; wide field | relative | Minimal | 30°C | mid-log |
| CHO | Chong et al. 2015 [8] | GFP-microscopy | live cells; confocal | relative | Minimal | 30°C | mid-log |
| YOF | Yofe et al. 2016 [61] | GFP-microscopy | N-terminal GFP; live cells; confocal | relative | Minimal | 30°C | mid-log |
| NEW | Newman et al. 2006 [37] | GFP-flow cytometry | live cells | relative | YPD | 30°C | mid-log |
| LEE | Lee et al. 2007 [29] | GFP-flow cytometry | live cells | relative | YPD | 30°C | mid-log |
| DAV | Davidson et al. 2011 [11] | GFP-flow cytometry | live cells | relative | YPD | 30°C | mid-log |
| GHA | Ghaemmaghami et al. 2003 [19] | TAP-immunoblot | SDS extract; immunoblot with internal standard | absolute | YPD | 30°C | mid-log |

For the purposes of our analysis, we consider the yeast proteome to consist of 5858 proteins [*Saccharomyces* Genome Database, www.yeastgenome.org, accessed October 28, 2016], encoded by 5157 verified ORFs and 701 uncharacterized ORFs. We excluded 746 dubious ORFs, as defined in the *Saccharomyces* Genome Database, from our analysis. Although some had peptides detected by mass spectrometry as annotated in the PeptideAtlas (http://www.peptideatlas.org) and Global Protein Machine (GPM, http://www.thegpm.org) databases, only 6 had good evidence for expression as defined by GPM. Proteins encoded by transposable elements, although readily detected, are not included in our analysis because most do not map to a unique ORF. Abundance data were called out of each of the 19 datasets using the 5858 protein ORFeome (Table S1).

### Data Transformation and Assessing Correlation

The natural logarithm was taken for each data set, since this is approximately normally distributed and thus suitable for linear regression analyses. All analyses and calculations were performed on natural log transformed data, unless specified otherwise. The Pearson correlation coefficient (r) was used for all correlation analyses in our study.

### Normalization of Arbitrary Unit Abundance Values

Mode shift normalization was applied to all studies that measured relative protein abundance and reported values in arbitrary units (Table 1). Each study that required normalization was natural log transformed and divided into 50 bins of equal abundance range. The median value of the bin with the greatest number of observations (values reported) was defined as the mode of the distribution. A scalar value was applied to each study to shift the mode to an arbitrarily chosen value of 100 arbitrary units. Mode shift normalized values were used for the remainder of the analysis.

For comparison to the mode shift normalization, studies were also quantile normalized and center log ratio transformed. For quantile normalization, proteins with a reported measurement from every study were retained for analysis, and quantile normalization was performed as described (Qiu et al. 2013). To normalize data sets by the center log ratio transformation method, arbitrary abundance measurements for each protein from each study were divided by the geometric mean and log10 transformed: *centerlog ratio = log* _*10*_*(Xi / geometric mean (X))*.

### Converting Protein Abundance From Arbitrary Units to Molecules per Cell

Mean protein abundance for each ORF was calculated for the four mass spectrometry-based studies reporting absolute protein abundance (Lu et al. 2007; Peng et al. 2012; Kulak et al. 2014; Lawless et al. 2016). We used the mean values as our calibration set (Table S1), as all four studies measured untagged proteins and reported protein abundance in molecules per cell. The mean abundance for each protein was natural log transformed and a least-squares linear regression was fitted between the calibration set and the natural log transformed mean mode-shifted arbitrary units, resulting in the following equation:

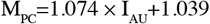

Protein molecules per cell (M_PC_) were then estimated from arbitrary intensity units (I_AU_) by applying the linear regression model to each individual dataset reported in arbitrary units.

### Calculating Coefficients of Variation

For each ORF, the coefficient of variation (CV) was calculated by:

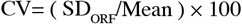

The CV was calculated for each ORF when at least two measurements were reported.

### Outlier Detection between GFP and Mass Spectrometry Studies

The mean abundance value for each ORF was calculated for MS-and GFP-based studies. A least squares linear regression model was fitted between these variables. This regression was used to identify point leverage by calculating hat values, outliers by calculating studentized residuals, and influential observations by calculating Cook’s distance. Outliers were defined as observations with studentized residuals greater than 2 or less than −2.

### Gene Ontology Term Enrichment

GO term analysis was performed using the GO term finder tool (http://go.princeton.edu/) using a P-value cutoff of 0.01 and applying Bonferroni correction, querying biological process or component enrichment for each gene set. After removing high frequency terms (>10% of background gene set), GO term enrichment results were further processed with REViGO (Supek et al. 2011) using the “Medium (0.7)” term similarity filter and simRel score as the semantic similarity measure.

### Spatial Analysis of Functional Enrichment (SAFE)

Functional annotation of protein abundance measurements on available genetic similarity networks constructed by Constanzo et al. (2016) was performed as previously described (Baryshnikova 2016), using Cytoscape v3.4.0 (Cline et al. 2007; Shannon et al. 2003).

All statistical analysis, data manipulation, and data visualization was performed in R (https://www.r-project.org).

## Results

### Comparisons of global quantifications of the yeast proteome

With 19 global quantitative studies of the yeast proteome (Ghaemmaghami et al. 2003; Newman et al. 2006; Lee et al. 2007; Lu et al. 2007; de Godoy et al. 2008; Davidson et al. 2011; Lee et al. 2011; Thakur et al. 2011; Nagaraj et al. 2012; Peng et al. 2012; Tkach et al. 2012; Breker et al. 2013; Denervaud et al. 2013; Mazumder et al. 2013; Webb et al. 2013; Kulak et al. 2014; Chong et al. 2015; Lawless et al. 2016; Yofe et al. 2016), 14 of which are reported in arbitrary units, we sought to derive absolute protein molecules per cell for the proteome for each data set and analyze the resulting data. We extracted the raw protein abundance values from the 19 datasets (Table S1) for the 5858 proteins in the yeast proteome, and compared the values (absolute abundance or arbitrary units) from each study with one another, resulting in 171 pairwise correlation plots (Figure 1). The studies agree well with one another, with Pearson correlation coefficients (r) ranging from 0.46 – 0.96. Notably, all studies with abundance measurements derived from GFP fluorescence intensity correlate better with one another than they correlate with the TAP-immunoblot or mass spectrometry-based studies.

**Figure 1.**
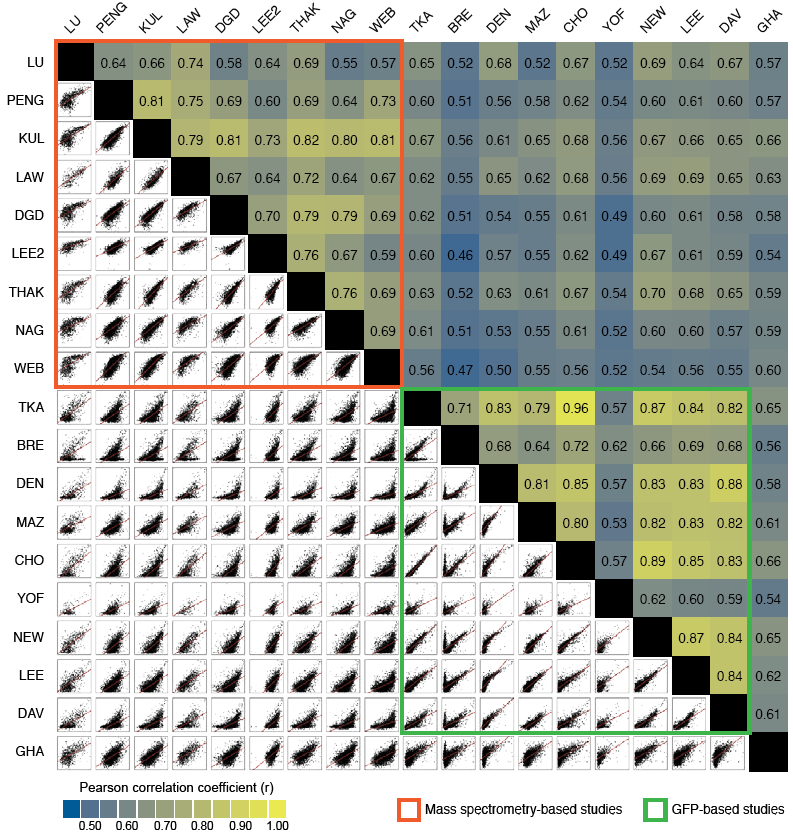
Scatterplot matrix of pairwise comparisons between protein abundance studies. Protein abundance measurements from 19 studies were natural log transformed and each pairwise combination was plotted as a scatterplot (bottom left). The least squares best fit for each pairwise comparison is shown (red line). The corresponding Pearson correlation coefficient (r) for each pairwise comparison is shown (top right) and shaded according to the strength of correlation. Mass spectrometry-based studies are indicated in orange, and GFP-based studies are indicated in green. Each abundance study is indicated by a letter code as described in Table 1.

### Protein copy number in S. cerevisiae

The most intuitive expression of protein abundance is molecules per cell. In order to convert all 19 datasets to a common scale of molecules per cell we had to first normalize the datasets and then apply a conversion factor to those data not expressed in molecules per cell. The experimental design, data acquisition and processing for the different global proteome analyses differ between studies. Moreover, the method for calculating GFP fluorescence intensity differs between high-throughput microscopic analyses. For example, (Chong et al. 2015) measured mean GFP intensity whereas (Tkach et al. 2012) calculated the integrated GFP intensity. As a result, protein abundance is reported on drastically different scales (Figure S1A). We tested three different methods to normalize all the data reported in arbitrary units (mode shifting, quantile normalization, and center log ratio transformation). The results of all three methods of normalization correlate very highly with one another (r = 0.96 – 0.97) indicating that the protein abundance values we calculate are largely independent of the specific normalization technique applied (Figure S1B). We also considered a normalization scheme where each protein is quantified relative to all other proteins in the dataset, as was done in PaxDb (Wang et al. 2012; Wang et al. 2015). While this relative expression of abundance (parts per million) has the advantage of being independent of cell size and sample volume, it makes comparison between different datasets difficult if the datasets measure different numbers of proteins. Thus, the parts per million normalization alters the pairwise correlations between datasets (Figure S2). By contrast, normalization by mode-shifting or center log ratio transformation allows comparison between datasets by expressing them on a common scale (Figure S1) and preserves the correlations that are evident in the raw data (Figure S2). Normalization by mode-shifting or center log ratio transformation also allows us to retain proteins whose abundance is not reported in all data sets in our aggregate analysis, thereby affording the greatest possible proteome coverage.

Currently five protein abundance data sets are reported in molecules per cell, four of which are mass spectrometry–based studies and one of which used an immunoblotting approach (Lu et al. 2007; Peng et al. 2012; Kulak et al. 2014; Lawless et al. 2016; Ghaemmaghami et al. 2003). The four mass spectrometry studies correlate well with one another (r = 0.64 to 0.81; Figure 1) and all measure native unaltered proteins, and so we reasoned that they could be used to generate a conversion from relative protein abundance in arbitrary units, to molecules per cell. We used the mean of these four data sets as a calibration dataset to convert every other dataset to molecules per cell. Although it is difficult to discern the accuracy of the protein abundance values in the calibration dataset, we find that the median difference between the calibration dataset values and the protein abundance values reported for 38 proteins in two small scale, internally calibrated studies (Picotti et al. 2009; Thomson et al. 2011), was 1.6-fold (Table S2), suggesting that protein abundance measurements from large scale studies are similar to those from smaller scale studies.

To convert all datasets to molecules per cell, a least-squares linear regression between the natural log transformed calibration dataset (reported in molecules per cell) and the natural log transformed mode-shifted or center log transformed studies (reported in arbitrary units) was generated. The correlation between the calibration dataset and the aggregate mode-shifted dataset was similar to the center log transformed dataset (Figure S1C) but had a lower sum of standardized residuals, so we proceeded with normalization by mode-shifting. Conversion of GFP measurements to molecules per cell resulted in a unified dataset covering 97% of the yeast proteome (Table S3). Of the 5858 protein proteome, only 156 proteins were not detected in any study (Table S4). The 156 proteins are enriched for uncharacterized ORFs (hypergeometric p = 6.9 × 10^−81^) and for genes involved in proton transport and glucose import (p = 5.9 × 10^−5^ and p = 0.0080, respectively). 353 proteins were detected in only a single study.

In general, there is agreement in the molecules per cell for each protein among the data sets analyzed in our study, with protein abundances ranging from 5 to 1.3 × 10^6^ molecules per cell (Figure 2A and Table S3). Notably, the mass spectrometry-based analysis by Kulak et al. exhibits the greatest sensitivity, reportedly capable of measuring less than 50 molecules per cell, and has the greatest detection range (Kulak et al. 2014). Many studies only provide values of protein copy number above ~2000 molecules per cell (Figures 2B and 3A). In particular, the GFP fluorescence–based studies tend to have a limited ability to detect low abundance proteins, likely because cellular autofluorescence presents a large obstacle to measuring the levels of low abundance proteins. In fact, one GFP-based study removed proteins whose fluorescence was close to background from their analysis (Chong et al. 2015), and in all GFP-based studies there are few values reported below 1000 molecules per cell (1794 values of 32318 reported).

**Figure 2.**
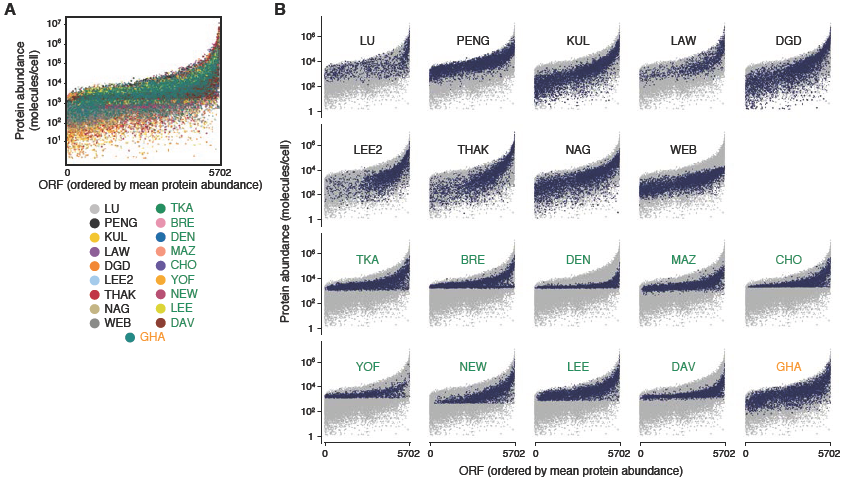
Protein abundance in nineteen data sets, in absolute molecules per cell. (A) The nineteen protein abundance data sets were normalized and abundance measurements were converted to molecules per cell and plotted. The mean protein abundance value, in molecules per cell, was calculated for each protein, and the proteins were ordered by increasing mean abundance on the x-axis. (B) Detected and quantified proteins from each study are highlighted (blue) and plotted with the abundance measurements from all nineteen data sets (grey). Proteins are ordered by increasing mean abundance along the x-axis. Letter codes are as in Table 1. Mass spectrometry-based studies are indicated in black text, GFP-based studies in green, and the TAP-immunoblot study in orange.

Genetic interaction networks have been extensively characterized in yeast, mapping genes and pathways into functional modules (Costanzo et al. 2016). We used spatial analysis of functional enrichment (SAFE) (Baryshnikova 2016) to identify the regions of the genetic interaction similarity network (Costanzo et al. 2016) that are enriched for high and low abundance proteins in our normalized protein abundance dataset (Figure 3B). We found high abundance proteins were specifically overrepresented in network regions associated with cell polarity and morphogenesis, and with ribosome biogenesis (Figure 3B, orange). Low abundance proteins were overrepresented in the region associated with DNA replication and repair (Figure 3B, teal).

**Figure 3.**
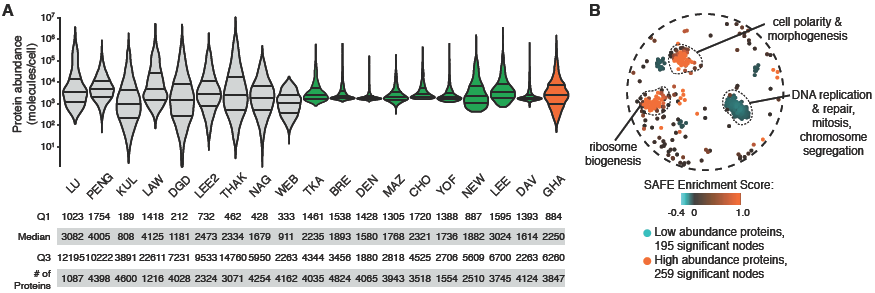
Distributions of protein abundance and functional enrichment. (A) The distribution of yeast protein abundance, as measured in each independent study in molecules per cell, is plotted, with the first quartile (Q1), median, and third quartile (Q3) indicated by horizontal bars. The areas of the violin plots are scaled proportionally to the number of observations. Mass spectrometry–, GFP–, and TAP immunblot–based studies are coloured in grey, green, and orange, respectively. The number of proteins detected and quantified by each study is also indicated. (B) SAFE annotation of the yeast genetic interaction similarity network (Costanzo et al. 2016) with protein abundance data. The protein abundance enrichment landscape is shown. Coloured nodes represent the centers of local neighborhoods enriched for high or low abundance proteins, shaded according to the log enrichment score. The outlines of the GO-based functional domains of the network where protein abundance enrichment is concentrated are shown.

GO-term enrichment analysis yielded results consistent with SAFE analysis. The decile comprising the least abundant proteins was enriched for DNA recombination (p = 2.7 × 10^−3^) and protein ubiquitination (p = 1.3 × 10^−4^), perhaps reflecting a limited requirement for these processes during unperturbed cell proliferation. The most highly expressed proteins tended to be proteins involved in translation in the cytoplasm (p = 3 × 10^−122^) and related processes, consistent with the key role of protein biosynthetic capacity in cell growth and division (Warner 1999; Volarevic et al. 2000; Jorgensen et al. 2002; Bernstein and Baserga 2004; Yu et al. 2006; Bjorklund et al. 2006; Teng et al. 2013).

### Variance in protein abundance measurements

Since each of the 19 studies in our analysis independently measured protein concentration using different methodologies and analyses, we explored the variation in reported values for each ORF among the 19 experiments. We calculated the coefficient of variation (CV; standard deviation / mean, expressed as a percentage) across the yeast proteome. The greatest median CVs were exhibited by proteins with the lowest and the highest abundance (Figure 4). While many factors can contribute to variation in the data, (e.g., experimental design, differences in media composition and acquisition of data) variation is the least for protein abundances between ~1000 and ~2000 molecules per cell, and the lowest median CV values are reported for abundances ranging from 1151 – 1337 molecules per cell.

**Figure 4.**
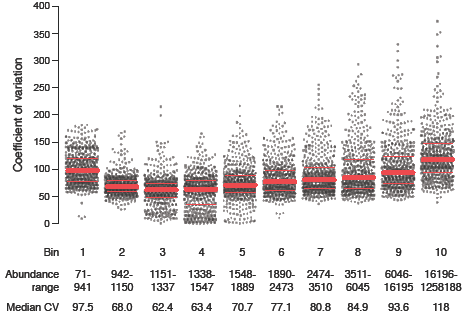
Variability of each protein abundance measurement. Proteins were ordered by increasing mean abundance and then binned into deciles. The coefficient of variation was calculated for each protein and plotted. The protein abundance levels associated with each bin are indicated below the scatter plot, as is the median CV for each bin. The red lines indicate the third quartile, the median, and the first quartile for each bin.

### Comparison of RNA expression and translation rates to protein abundance

For most proteins, expression appears to be tightly regulated to maintain levels within an appropriate range of copies per cell. To determine the contribution of RNA expression to steady-state protein levels, we compared protein copy number per cell to mRNA levels from three microarray and three RNA-seq datasets (Roth et al. 1998; Causton et al. 2001; Lipson et al. 2009; Nagalakshmi et al. 2008; Yassour et al. 2009). In general, our protein abundance values correlate with mRNA levels as measured by microarray (r = 0.60 – 0.68) and RNA-seq (r = 0.65 – 0.75) (Figure 5A). Higher correlations between mRNA and protein abundance have been reported (r = 0.66 - 0.82) (Futcher et al. 1999; Greenbaum et al. 2003; Franks et al. 2015) in studies using less comprehensive protein abundance datasets (2044 proteins at most), suggesting that a more complete view of the relationship between transcript and protein abundance could be obtained using our more comprehensive protein abundance dataset. In addition to capturing a large fraction of mRNA abundance variance, our protein abundance dataset correlates well with translation rates derived from ribosome profiling studies (McManus et al. 2014) (r = 0.71, Figure 5B).

**Figure 5.**
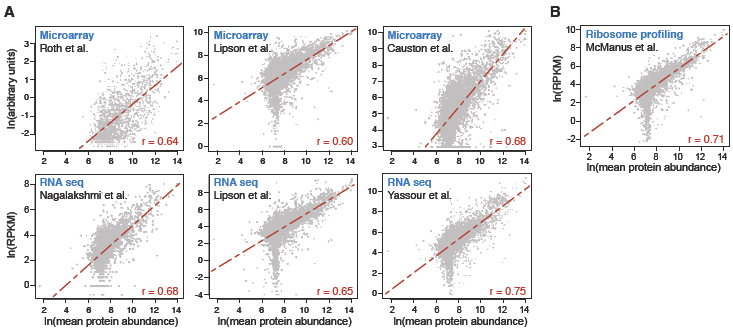
Comparison of protein abundance with mRNA levels. (A) Protein abundance (natural log of the mean number of molecules per cell) is compared with mRNA levels measured by microarray analyses from three independent studies (natural log of arbitrary units), and with mRNA levels measured by RNA sequencing analysis (natural log of reads per kilobase of transcript per million mapped reads (RPKM)). (B) Protein abundance (natural log of mean molecules per cell) is compared with ribosome footprint abundance (natural log of RPKM) from ribosome profiling analysis.

### Protein fusion tags have limited effect on native protein abundance

The yeast strains used to measure protein abundance and localization by GFP fluorescence all express proteins with C-terminal fusions to GFP (Huh et al. 2003), with the exception of the Yofe et al. study, which analyzed 1554 N-terminal GFP fusions (Yofe et al. 2016). C-terminal fusion to GFP sequences adds an extra 27 kDa to the native protein, alters the identity of the C-terminus, and changes the DNA sequence of the 3’ untranslated region of the gene. Evidence suggests that fusion to GFP has a limited effect on the intracellular localization of the proteome (Huh et al. 2003), although a fraction of the proteome is inaccessible with C-terminal tags (Yofe et al. 2016). The effect of fusion to GFP on protein abundance has yet to be examined systematically. We reasoned that proteins whose expression differs greatly between mass spec data sets (which measure native proteins) and GFP data sets are likely affected by the presence of the tag. We compared the mean ln abundance between mass spectrometry and GFP-based abundance studies. Since they correlate well (r = 0.68), we fitted a least squares linear regression model through the ln-transformed data. We used a studentized residuals threshold approach, defining proteins with studentized residuals greater than 2 and less than −2 as outliers (Figure 6 and Table S5). A total of 260 proteins were identified, with 107 proteins exhibiting greater abundance in the native protein state, ranging from 5-fold to 221-fold, compared to the GFP-tagged protein (Figure 6B, C). The 107 proteins were enriched for ribosome components (29 proteins; p = 1.1 × 10^−11^), suggesting that caution is warranted when tagging ribosome subunits. Sixty-two of the 107 proteins with reduced abundance when GFP-tagged have also been assessed as C-terminal fusions to the 21 kDa TAP-tag (Ghaemmaghami et al. 2003). Of the 62, 37 proteins also had reduced abundance (by at least 2-fold) when TAP-tagged, suggesting that these proteins, again enriched for ribosome components (14 proteins; p = 8.5 × 10^−7^), are either destabilized by the presence of any protein tag at the C-terminus, or require their native 3’ UTR for mRNA stability (Figure 6D). Interestingly, 26 of the 107 proteins with reduced abundance when C-terminal GFP-tagged were also assessed as N-terminal GFP fusions (Yofe et al. 2016). All but one had reduced abundance (by at least 2-fold) irrespective of the location of the GFP tag. Twenty-five proteins decreased in abundance when GFP-tagged but not when TAP tagged. These 25 proteins, which were not enriched for any GO process terms, could represent GFP-specific protein destabilization, or protein-specific issues with fluorescence detection (Waldo et al. 1999).

**Figure 6.**
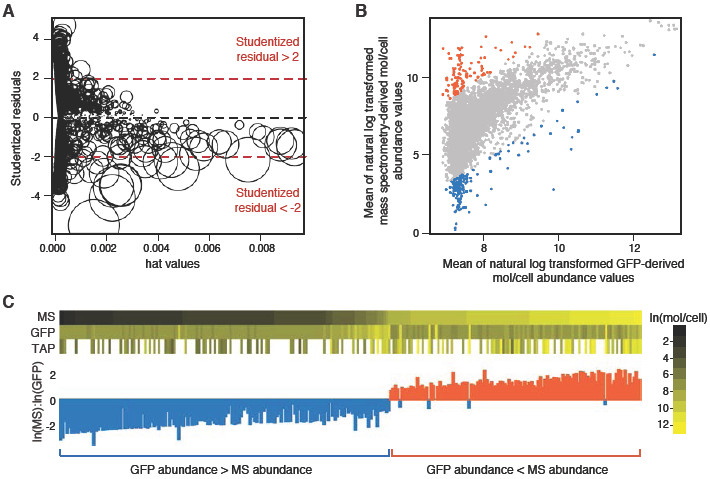
Identification of proteins whose expression is influenced by protein fusion tags. (A) The natural log transformed mean protein abundance from mass spectrometry-based studies is compared to the natural log transformed mean protein abundance from GFP-based studies and displayed as an influence plot to identify data points with undue influence over the fitted regression equation. Studentized residuals are plotted against leverage (hat values), and point size reflects Cook’s distance. (B) Means of mass spectrometry-derived abundance values are plotted against means of GFP-derived abundance values. Outliers with residuals >2 or <-2 in the influence plot are highlighted in red and blue, respectively. (C) Mean mass spectrometry (MS), TAP-immunoblot (TAP), and GFP protein abundance values for each identified outlier are compared. Proteins are ordered by increasing mass spectrometry abundance, with each bar representing a single protein. The ratio of the natural log of the MS abundance to the natural log of the GFP abundance is displayed for each outlier.

We also observed 153 outliers that had greater abundance in GFP studies than in mass spectrometric analyses, by 1.5-fold to as much as 2700-fold. While it is possible that a GFP-fusion tag could cause an increase in the abundance of a given protein, perhaps by increasing protein stability, one likely explanation is that many of these proteins are in low copy number per cell and therefore below the accurate detection limit for GFP fluorescence intensity (Figure 6C). A conservative estimate of the lower limit for accurate detection of GFP fluorescence intensity is the minimum number of molecules per cell reported in any GFP-based study, 370. Removing outliers with less than 370 molecules per cell as measured by mass spectrometric methods leaves only 31 proteins that increased in abundance when fused to GFP. Thus, although some specific changes in abundance occur for a small number of proteins upon adding additional sequences to the C-terminus, it appears that most yeast proteins (95% of the 5342 proteins measured with C-terminal GFP tags) can tolerate C-terminal tags without large changes in protein expression.

### Changes in protein abundance under environmental stresses

External stressors can perturb cellular processes and activate the environmental stress response, a mechanism for cells to protect themselves from fluctuating conditions in the environment (Gasch et al. 2000; Gasch and Werner-Washburne 2002). High-throughput (HTP) fluorescence microscopy and mass spectrometry have enabled large scale analyses of the proteome after exposure to diverse stresses, including quiescence, DNA replication stress conditions, oxidative stress, nitrogen starvation, reductive stress, and rapamycin treatment. Given that protein concentration directly influences cellular processes and function, we were interested in quantifying absolute protein molecules per cell and comparing changes in protein levels across studies investigating condition-dependent protein abundance changes. To simplify the comparisons, we focused on GFP-based studies, which are available for hydroxyurea, methyl methanesul-fonate, oxidative stress, reductive stress, nitrogen starvation, rapamycin treatment, and quiescence (Davidson et al. 2011; Tkach et al. 2012; Breker et al. 2013; Denervaud et al. 2013; Mazumder et al. 2013; Chong et al. 2015). Mass spectrometry datasets are available for diploid cells, heat shock, high salt, quiescence, and 13 different carbon sources (de Godoy et al. 2008; Nagaraj et al. 2012; Lee et al. 2011; Webb et al. 2013; Usaite et al. 2008; Paulo et al. 2015; Paulo et al. 2016), but are not considered here.

Since the majority of proteins do not change in abundance in any given stress condition, we normalized GFP intensities from each study by the mode-shifting method and applied the same linear regression used previously to convert arbitrary units to protein molecules per cell (Table S6). We applied a cut-off for changes in protein abundance, corresponding to either a two-fold increase or a two-fold decrease (Table S7). At this cut-off, which is more conservative than that used in most of the individual studies, 1250 of 4263 proteins assessed change in abundance in at least one condition: 580 proteins increase in abundance, and 744 proteins decrease in abundance. The magnitude of abundance changes spans a range of 60-fold for increases and 57-fold for decreases (Table S7; the Lee et al. dataset was excluded from analysis of abundance decreases as its inclusion results in maximum-fold decreases that greatly exceed the dynamic range of GFP fluorescence detection that is evident in Figure 3). Proteins that increased or decreased in abundance during stress tended to be of higher abundance in unperturbed cells than the proteome median (Figure S3).

Eighty-two percent of the abundance changes observed were specific to one or two conditions, suggesting significant stress-specific regulation. Two proteins, Hsp12 and Ynl134c, were the most universal stress responders, increasing in abundance in 9 of 11 perturbation datasets. Finally, we note that in the case of MMS treatment, where four datasets are available (Lee et al. 2007; Tkach et al. 2012; Denervaud et al. 2013; Mazumder et al. 2013), only a single protein (Yml131w) has a statistically supported abundance change greater than 2-fold when the four datasets are compared. Since the conditions of growth, treatment, image acquisition, and image analysis differ between studies, we suggest that use of standardized protocols will be the first step towards evaluating protein abundance changes during stress conditions.

## Discussion

Here we provide a comprehensive view of protein abundance in yeast by normalizing and combining 19 abundance datasets, collected by mass spectrometry (Lu et al. 2007; de Godoy et al. 2008; Lee et al. 2011; Thakur et al. 2011; Peng et al. 2012; Nagaraj et al. 2012; Kulak et al. 2014; Lawless et al. 2016), GFP fluorescence flow cytometry (Newman et al. 2006; Lee et al. 2007; Davidson et al. 2011), GFP fluorescence microscopy (Tkach et al. 2012; Breker et al. 2013; Denervaud et al. 2013; Mazumder et al. 2013; Chong et al. 2015; Yofe et al. 2016), and western blotting (Ghaemmaghami et al. 2003). Despite different experimental design, conditions, and methodologies for detection and analysis, protein abundance correlates well between the different studies, ranging from r = 0.46 – 0.96. Correlation is highest among the datasets collected using GFP fluorescence, however the detection range of GFP intensities is limited. Low abundance proteins are difficult to distinguish from cellular auto-fluorescence, and high abundance protein intensity measurements approach saturation of the GFP fluorescence detection. Our estimates of mean protein abundance could likely be improved if cellular autofluorescence levels were reported in the GFP intensity datasets such that low-confidence, low fluorescence values could be filtered. The mass spectrometry-based analyses provide the greatest sensitivity and dynamic range for protein measurements, reporting protein abundance measurements for the full range of abundance levels in the proteome. Collectively, our analysis suggests protein abundance in the yeast proteome ranges from zero to 1.3 × 10^6^ molecules per cell. Interestingly, 75% of yeast proteins quantified are present at between 1000 and 10 000 molecules per cell, indicating that it is rare for proteins to be present at very high or very low copy numbers.

Measurements of coefficients of variation reveal that while there is variance in abundance measurements across the entire range of abundance values, the greatest variation is exhibited at the abundance extremes. Measurement of low abundance proteins is confounded by detection and resolution limits of all but the most sensitive mass spectrometry-based approaches. Highly abundant proteins likely have greater variance for two reasons: (1) GFP-based quantification of highly expressed proteins underestimates the true value due to saturation, and (2) the correlation between MS-based studies is lesser for highly expressed proteins than it is on average.

Construction of the TAP and GFP collections involved tagging the 3’ end of each annotated ORF, at the chromosomal locus. Our data indicate that C-terminal tags have little effect on the abundance of most proteins, since 95% (5083 of 5343) of the proteins measured showed no large change in abundance when tagged. Only 515 proteins are not represented in the C-terminal GFP datasets, and of these 156 were not detected by any method, and so it is unlikely tagging specifically destabilized these proteins. We infer that at most an additional 359 proteins could be affected by tagging, leaving 89% of the yeast proteome unaffected by C-terminal tagging. Thus, the 515 proteins absent from existing datasets are unlikely to affect the general conclusion that the yeast proteome can tolerate C-terminal tags well, without large effects on protein expression levels.

The yeast proteome is dramatically remodeled in response to stress (Tkach et al. 2012; Breker et al. 2013; Denervaud et al. 2013; Mazumder et al. 2013; Chong et al. 2015; Breker et al. 2014; Koh et al. 2015). Upon aggregating all condition-dependent studies and providing absolute protein abundance values in stress conditions we found that 1250 yeast proteins experienced an abundance change of at least 2-fold (Table S7). This is almost certainly an underestimate. Most stress condition studies rely on the GFP collection, which covers only 70% of the proteome. Further, the number of environmental conditions that have been assessed to date is considerably fewer than those for which mRNA abundance data is available. Therefore, it is perhaps surprising that only 234 proteins in this analysis (4% of the proteins assessed as GFP fusions) were upregulated or downregulated in even three stress conditions. This contrasts with the almost 900 core stress response genes identified through microarray analyses (Gasch et al. 2000; Hughes et al. 2000; Causton et al. 2001). What could account for this apparent discrepancy? First, the half-life of a typical mRNA (~32 minutes) (Geisberg et al. 2014) is short compared to the typical protein (~ 43 minutes) (Belle et al. 2006), and so it might be expected that mRNA levels would show a more rapid response to stress than would protein levels. Secondly, diverse post-transcriptional regulation modes can be brought to bear on protein function, including regulation of translation, protein degradation, protein modification, and intracellular localization changes so protein function needn’t be altered at the level of abundance alone. Finally, the environmental stress response at the mRNA level was typically defined by clustering analysis, rather than by-fold mRNA abundance changes, and so many core transcriptional responses are below the 2-fold cutoff that we applied to the protein abundance data. Protein abundance directly influences cellular processes and phenotypes. The plasticity of the proteome in stress conditions has been extensively investigated in yeast. We unified the available data and report protein abundance in a single common unit of molecules per cell, in both unperturbed cells and in response to stress, providing a useful resource for further analysis of the dynamic regulation of the proteome.

**Figure S1.**
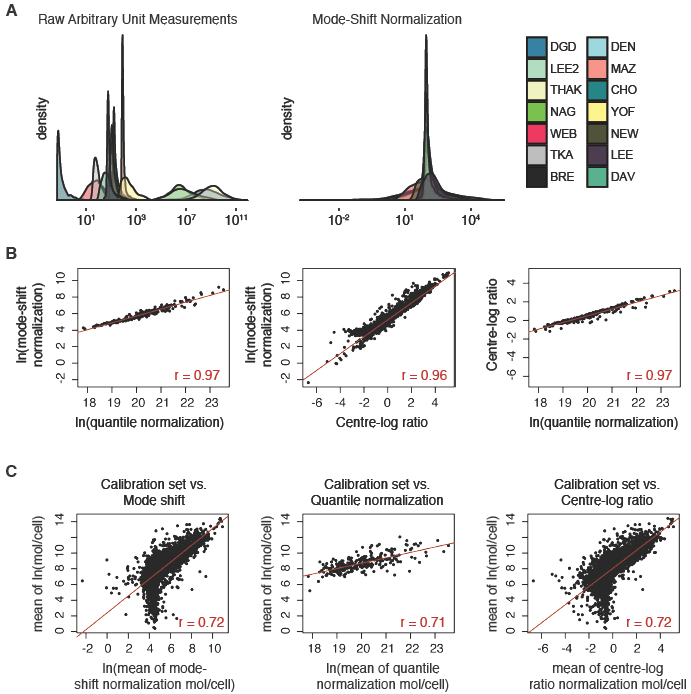
Normalization methods and comparisons to the calibration abundance data set. (A) Raw protein abundance measurements from studies reporting arbitrary units (left) were mode-shift normalized (right). (B) Protein abundance measurements were normalized using mode-shift, quantile, or centre-log ratio normalization methods. The mean abundance for each protein was calculated following each normalization, and each was compared to the others. (C) Each normalization method was compared with the mean abundance from the calibration data set. Pearson correlation coefficients (r) are indicated in each plot.

**Figure S2.**
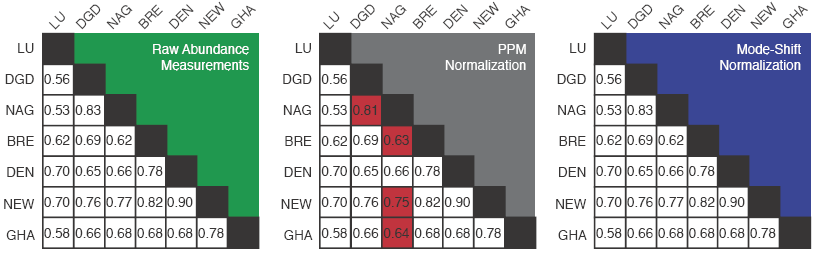
Comparison of parts per million and mode-shift normalization methods. Pearson correlation coefficients were calculated for each pairwise comparison of the seven studies indicated. Correlation coefficients were calculated for the raw abundance measurements (left), parts per million normalized datasets (middle), or mode-shifted datasets (right). Boxes shaded in red indicate correlations that are not equivalent to correlations among the original raw datasets.

**Figure S3.**
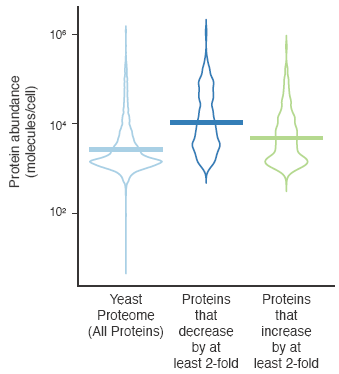
Abundance distributions of the proteome and proteins that change in stress. The protein abundance distribution in molecules per cell is presented as a violin plot for all proteins in the proteome, for proteins that decrease in abundance by 2-fold in at least one study, and for proteins that increase in abundance by 2-fold in at least one study. The horizontal bars represent the medians, and violins are scaled such that all have the same area.

## Acknowledgments

This work was supported by the Canadian Cancer Society Research Institute (Impact grant 702310 to GWB), an Ontario Government Scholarship and a Natural Sciences and Engineering Research Council of Canada CGS-M award (to BH), and a Lewis-Sigler Fellowship (to AB). We thank Helena Friesen, Tina Sing, and Xanita Saayman for helpful discussions and careful reading of the manuscript.

## Author contributions

BH, AB, and GWB designed and performed the analysis. BH and GWB wrote the manuscript.

